# Complement C3 reduces apoptosis in human cardiomyocytes

**DOI:** 10.1101/2023.05.01.538962

**Authors:** Zhou Fang, Xiang Li, Fajun Yang, Alus Michael Xiaoli, Ming Zhang

## Abstract

Complement C3 is a key factor in complement system. Our recently animal study found that C3 may regulate myocardial apoptosis through the intrinsic apoptosis pathway. The current work investigated if C3 regulation of apoptosis occurred in human cardiomyocytes. Our results showed that incubation of exogenous C3 reduced apoptosis in a cell culture system of human cardiomyocytes which did not inherently express C3. In addition, C3 inhibited intrinsic apoptosis pathway in a cell-free apoptosis system. Furthermore, pro-C3 was found to bind with an apoptotic factor, pro-caspase 3, in a cell-free system. Thus, we presented firsthand evidence that exogenous C3 is readily reduce apoptosis in human cardiomyocytes via interaction with the intrinsic apoptotic pathway.

## 1. Introduction

The complement system is a critical component of both the innate and adaptive immune systems that augments the function of antibodies and phagocytes (Geisbrecht et al., 2022). Complement activation can be initiated through the classical, the alternative, and the lectin pathways (M. C. Carroll and Isenman, 2012; M. V. Carroll and Sim, 2011; Cheng et al., 2020; Dunkelberger and Song, 2010; Gorsuch et al., 2012; Ricklin and Lambris, 2013; Sacks and Zhou, 2012; Sarma and Ward, 2011; Trouw and Daha, 2011). Complement C3 is a central factor in the three activation pathways (M. C. Carroll and Isenman, 2012; M. V. Carroll and Sim, 2011; Cheng et al., 2020; Dunkelberger and Song, 2010; Gorsuch et al., 2012; Ricklin and Lambris, 2013; Sacks and Zhou, 2012; Sarma and Ward, 2011; Trouw and Daha, 2011). The three pathways converge at the key central factor, C3, and are followed by a common cascade (M. C. Carroll, 1998): C3 convertases yield C3a and additional C3b, the latter forming C5a convertases that cleave C5 to C5a and C5b. C5b initiates membrane attack complex (MAC) formation (C5b-C9) (Harboe and Mollnes, 2008; Ricklin et al., 2010).

C3 is mainly synthesized by hepatocytes (Li et al., 2021), starting as pro-C3, which is a single polypeptide chain. Pro-C3 was proteolytically processed to two chains, α-chain (120kDa) and β-chain (75kDa) (Nishida et al., 2006), which are linked by disulfate bonds and occurs intracellularly (Brade et al., 1977). After its processing, the double-chain C3 is secreted into the circulation (Bokisch et al., 1975; Fearon and Wong, 1983; Fontaine and Rivat, 1979; Minta et al., 1977; Parkes et al., 1981). Spontaneous hydrolysis of C3, which takes place in plasma (Fromell et al., 2020), generates its hydrolytic product C3(H_2_O). Further enzymatic cleavage of C3 alpha-chain, which may occur intracellularly and extracellularly (Liszewski et al., 2013), generates fragments of the overall molecule, termed C3a, C3b, C3c, C3d,g, C3d and C3g (Cheng et al., 2020).

We recently found that in a murine ischemia/reperfusion (IR) injury model, C3 plays a significant role in myocardial apoptosis possibly through interaction with factor(s) in the intrinsic apoptosis pathway, e.g. cytochrome *c* (Cyt *c*) (Fang et al., 2023). It is still unknown if exogenous C3 can affect myocardial apoptosis. In the current study, we investigated if exogenous C3 can affect apoptosis in human cardiomyocytes and explore relevant molecular mechanism.

## 2. Materials and Methods

### 2.1. Cell lines, Abs, molecular cloning construct, and proteins

AC16 human cardiomyocyte cell line (Millipore, MA) was grown in DMEM:F12 with 10% heat-inactivated FBS, 1% penicillin and 1% L-Glutamine. 293 T cell was grown in DMEM with 10% heat-inactivated FBS and 1% penicillin. Both cell lines were maintained at 37 °C in 5% CO_2_.

Rabbit anti-human C3c (F0201) polyclonal Ab was obtained from Agilent Dako (Santa Clara, CA). Goat anti-human C3 (A213) polyclonal Ab was obtained from Complement Technologies (Tyler, TX). Rabbit anti-Cyt *c* polyclonal Ab was obtained from Cell Signaling (Danvers, MA). Rabbit anti-caspase 3 polyclonal Ab was obtained from Santa Cruz Biotechnology (Dallas, TX). The full length human complement C3 expression ORF clone, whose C3 gene cDNA ORF clone sequences were retrieved from the NCBI Reference Sequence Database and the vector is pcDNA 3.1^+^/C-(K)DYK, was obtained from GenScript (Piscataway, NJ) (Supplementary Fig. 2).

### 2.2. qPCR for C3’s mRNA expression

C3 mRNA expression were studied by qPCR, and a positive control which internally expressed C3 was included by using Huh 7 human liver cell line (de Bruijn and Fey, 1985).

### 2.3. H2O2 induced apoptosis in human cardiomyocytes

Hydrogen peroxide (H_2_O_2_, Sigma-Aldrich, MO) induced apoptosis was performed in confluent AC16 cells. Cells were incubated with H_2_O_2_ for 60 minutes and then washed with PBS. Cells were then incubated in the presence or absence of purified human C3 for 3 hours. Apoptotic cell death was detected by FACS analyses of Annexin-V and propidium iodide staining at Washington University as reported previously (Kulkarni et al., 2019).

### 2.4. Cell Free Apoptosis Analyses

A cell free apoptosis system was employed (McCoy et al., 2013; Nutt et al., 2005). Cytosolic fractions from xenopus oocytes extracts were prepared and graciously supplied by Dr. Leta Nutt at St. Jude Hospital (McCoy et al., 2013; Nutt et al., 2005). Purified C3 was pre-incubated with either purified Cyt *c* (Sigma, MO) or a purified cytosolic fraction for 1 hour, followed by the either addition of purified cytosol from xenopus or purified Cyt *c*, and incubation for one hour. Apoptosis activities were detected using substrate from the Caspase-3 Glo Apoptosis kit (Promega, WI).

### 2.5. Pull-down assay to detect apoptotic factor(s) interacted with C3

#### 2.5.1. Transfection and overexpression of human C3 in 293 T cell line

293 T cells were split and placed in a 10 cm dish 4 hours before transfection. The DNA/CaPO4 mix for transfection was prepared as following: 10 μg of total DNA, 62 μl of 2M CaCl_2_, with additional ddH_2_O up to 500 μl total volume. This mixture was added to 500μl of 2X HBS at RT. DNA/CaCl2/HBS mixture was sat at RT for 25 minutes. Then the mixture was mixed again, following by sprinkling the entire mixture over the plate of cells. The cells were cultured in the medium with the transfection mixture for 36 hours to 48 hours, following by the collection of the cells.

#### 2.5.2. Apoptotic cell lysate preparation

To prepare apoptotic cell lysate of human cardiomyocytes, AC16 cells were incubated with 500 μM H_2_O_2_ (Sigma-Aldrich (St. Louis, MO)) for 30 minutes and then washed with PBS. The cells were collected. Cell lysate was prepared by resuspending cell pellets in IP buffer (0.2% NP-40, 10% Glycerol, 10mM Tris-HCl at pH 8.0, 100mM KCl) with 1% Halt™ Protease Inhibitor Cocktail (100X) (78430, Thermo Scientific) and incubating for 10minutes on ice. The resulting lysate were then centrifuged for 10 minutes at 8,000g and supernatants collected, aliquoted, and stored at –80°C. The same procedure for cell lysate was applied for the 273 T cells.

#### 2.5.3. Identification of apoptotic factor(s) binding to C3

This pull-down method is based on the binding affinity between antigen and antibody and the subsequent isolation of the capture/target complex by precipitation (Brymora et al., 2004; Dong and Li, 2018; Prelich, 2012). The anti-Flag antibody precoated beads were incubated with transfected 293 T cell lysate for 3 hours at 4°C. After incubation, the beads were collected and washed once with the IP buffer. After wash, the beads were incubated with AC16 cell lysate, which was treated with H_2_O_2_ for 3 hours at 4°C. The beads were washed 4 times after incubation, following by the elution the protein from the beads. The proteins in the protein complex were identified by the Western Blot.

### 2.6. Statistical analysis

Statistical analyses were performed using IBM SPSS Software version 20 (IBM Corp., NY). For animal studies, an independent *t*-test with two tails and unequal variances was used to determine the statistical significance of differences between the results of experimental and control groups. Descriptive data were summarized as mean ± standard error of mean.

## 3. Results

### 3.1. Reduction of apoptosis in AC16 cardiomyocytes by exogenous C3

We recently found that C3 block myocardial apoptosis in a murine heart IR model (Fang et al., 2023). To study if C3 has the same anti-apoptotic effect in human cardiomyocytes, we first analyzed if AC16 cells express C3 internally. C3 mRNA expression were studied by qPCR, and a positive control which internally expressed C3 was included (a human liver cell line, Huh 7) (de Bruijn and Fey, 1985). High levels of C3 expression were detected in Huh 7 cells, but AC16 cells did not have detectable C3 mRNA (Fig. 1a). Then, we induced apoptosis in AC16 cells by H_2_O_2_ treatment as reported (Xiang et al., 2016). When AC16 cells were incubated with high dose of H_2_O_2_ for 60 min, they underwent apoptosis as expected (Fig. 1b). However, addition of C3 into the cell culture significantly decreased apoptosis of AC16 cells (Fig. 1b). Therefore, although AC16 cells do not express C3, exposure of the exogenous C3 could reduce apoptosis in these cells under oxidative stress condition.

**Fig. 1.**
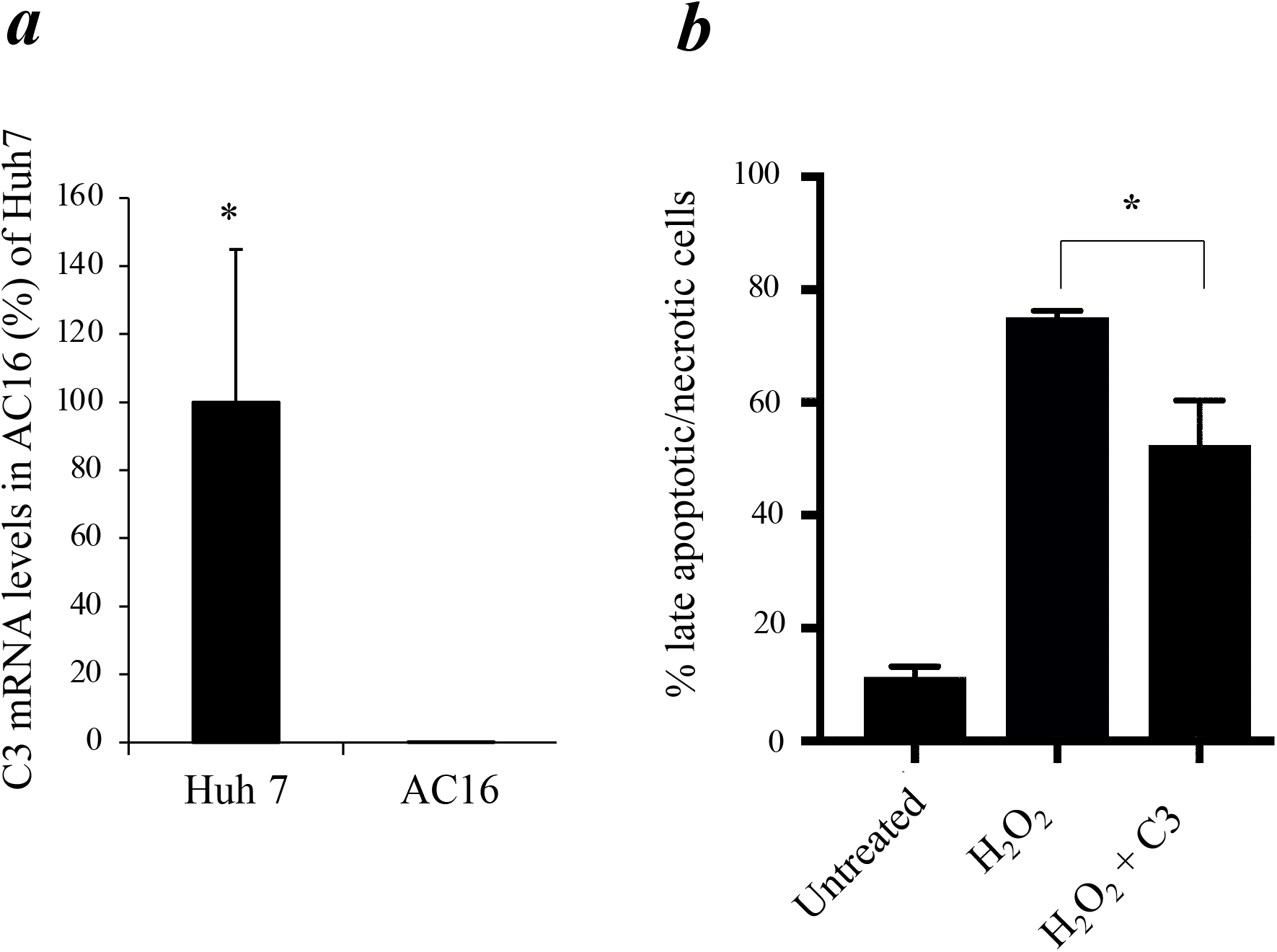
Exposure to exogenous C3 reduced apoptosis in cardiomyocytes. *a)*. C3 mRNA levels in AC16 cardiomyocytes were measured by qPCR. Huh 7 human liver cell line was used as a positive control of C3 expression. Error bars indicate SEM; *P<0.05. *b)*AC16 cells were incubated with H_2_O_2_ for 60 minutes and then washed with PBS. Cells were then incubated in the presence or absence of purified C3 for 3 hours. Apoptotic cell death was detected by FACS analyses of Annexin V-Propidium Iodide staining. *indicates P < 0.05.

### 3.2. Inhibition of the intrinsic apoptosis pathway by C3 in a cell-free system

Our recent study found that Cyt *c* was present in the C3-containing protein complexes of hearts in a murine IR injury model, suggesting that C3 may influence apoptosis by interacting with Cyt *c* in the intrinsic apoptotic pathway (Fang et al., 2023). Basic research have suggested that in the intrinsic apoptosis pathway, Cyt *c* released from mitochondria binds to Apaf-1 to form the apoptosome by recruiting caspase-9 (Liu et al., 1996; Zou et al., 1997), which further activates downstream caspases-3 and -7, leading to apoptosis (Konstantinidis et al., 2012).

To investigate the mechanism underlying C3’s interaction with Cyt *c* in the intrinsic apoptosis pathway, we employed a cell free apoptosis system (McCoy et al., 2013; Nutt et al., 2005). As outlined in Supplementary Fig. 1, we carried out experimental approaches as following: 1) pre-incubation of human C3 with Cyt *c*, followed by addition of the cytosolic fraction without mitochondria (containing factors of intrinsic apoptosis pathway except endogenous Cyt *c*); 2) pre-incubation of C3 with purified cytosol without mitochondria, followed by addition of Cyt *c*. Our hypotheses are: (i), if C3 binds Cyt *c* directly, Approach #1 will show reduction of apoptosis when C3 is pre-incubated with Cyt *c*, while Approach #2 will not show apoptosis reduction; (ii) On the other hand, if C3 binds to factor(s) downstream of Cyt *c* in the intrinsic apoptosis pathway, Approach #1 will not block apoptosis, while Approach #2 will block apoptosis when C3 is pre-incubated with purified cytosol followed by the addition of Cyt *c*.

In a control experiment without C3, incubation of Cyt *c* with purified cytosol induced apoptosis as expected, confirming this cell free system working as reported (McCoy et al., 2013; Nutt et al., 2005) (Fig. 2a and b, hatched bars). As a negative control, purified cytosol alone could not induce apoptosis without Cyt *c* (Fig. 2a and b, left hand bars). Pre-incubation of C3 with Cyt *c* did not block the Cyt *c* -mediated apoptosis (Fig. 2a, black bar). However, pre-incubation of C3 with purified cytosol blocked the Cyt *c* -mediated apoptosis (Fig. 2b, black bar). Thus, these data indicated that C3 does not bind Cyt *c* directly, but rather interacts with downstream apoptotic factor(s) in the cytosol in the intrinsic apoptosis pathway.

**Fig. 2.**
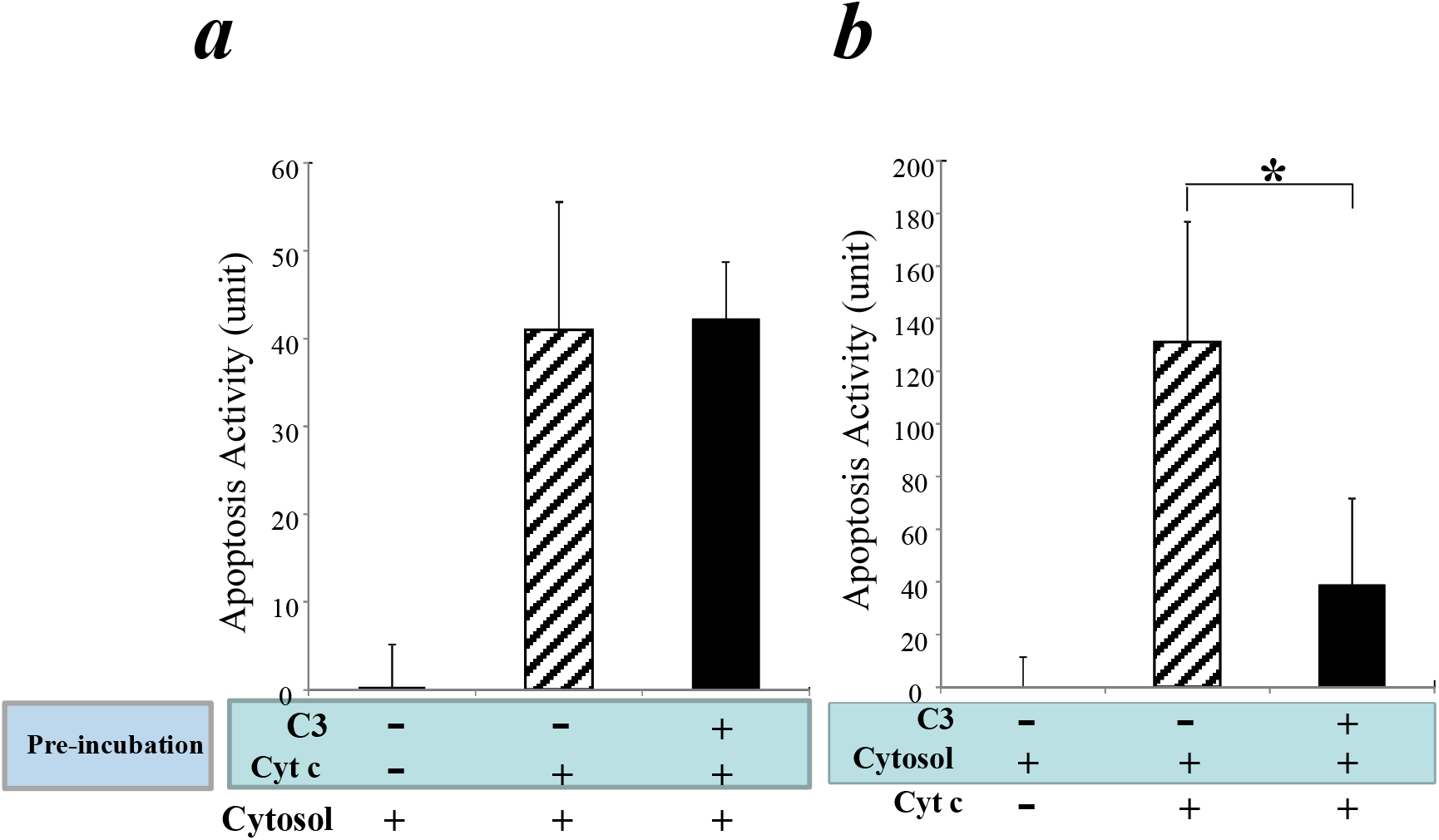
In a cell-free apoptosis system, C3 interacts with factor(s) downstream of Cyt *c* and blocks the Cyt *c* mediated apoptosis. a). Purified C3 was pre-incubated with purified Cyt *c* for one hour, followed by addition of purified cytosol from xenopus and incubation for one hour. Apoptosis activities were detected using substrate from the Caspase-3 Glo Apoptosis kit. b). Purified C3 was pre-incubated with purified cytosol for one hour, followed by addition of Cyt *c* and incubation for one hour. Apoptosis activities were detected as in (a). Error bars indicate SEM; *indicates P < 0.05.

### 3.3. Pro-C3 binding with pro-caspase 3 in a cell-free system

Previous animal studies by us and others have showed that under pathological conditions such as IR injury, circulation C3 was deposited in the ischemic myocardium flooded with oxygenated blood upon reperfusion (Charlagorla et al., 2013; Jordan et al., 2001), which could be a reason to reduce myocardial apoptosis (Fang et al., 2023). To further study the role of C3 in human cardiomyocyte apoptosis, we investigated if C3 is able to interact with the intracellular proteins related to apoptosis. We employed a method of antibody pull-down to identify potential intracellular targets bound to C3. To obtain a large amount of tagged-human C3 for pull-down assay, we transfected 293 T cells with a plasmid construct which overexpressed a full-length of single chain pro-C3 with a Flag-tag at its C-terminal (Flag-pro-C3; Supplementary Fig.2). The overexpressed Flag-pro-C3 was captured by beads pre-coated with anti-Flag antibody and served as the “bait” in the pull-down assay. The “prey” would be apoptosis related protein(s) in the cell lysate of AC16 cells treated with H_2_O_2_ which induced apoptosis in these cells (Xiang et al., 2016).

Pro-C3 proteins were readily detected in the pulldown complex using anti-Flag or anti-pro-C3 antibody (Fig. 3. Lane 3), demonstrating that transfection, overexpression and pulldown of Flag-pro-C3 proteins were successful. As a negative control, IP buffer was used instead of transfected 293 T cell lysate to incubate with the anti-Flag beads (Fig. 3. Lane 2). No C3 proteins were detected in the H2O2-treated AC16 cell lysate (Fig. 3. Lane 1), confirming that AC16 cells do not produce C3 inherently. When Flag-pro-C3 bound beads were incubated with the AC16 cell lysate to pull down proteins interacting with pro-C3, we could detect pro-caspase 3 in the pro-C3 binding complex (Fig. 3. Lane 3). Although both pro-caspase 3 and Cyt *c* are abundant in H2O2-treated AC16 cell lysates (Fig. 3. Lane 1), pro-C3 did not pull down any Cyt *c* proteins (Fig. 3. Lane 3), suggesting that Cyt *c* is not present in the pro-C3 binding complex.

**Fig. 3.**
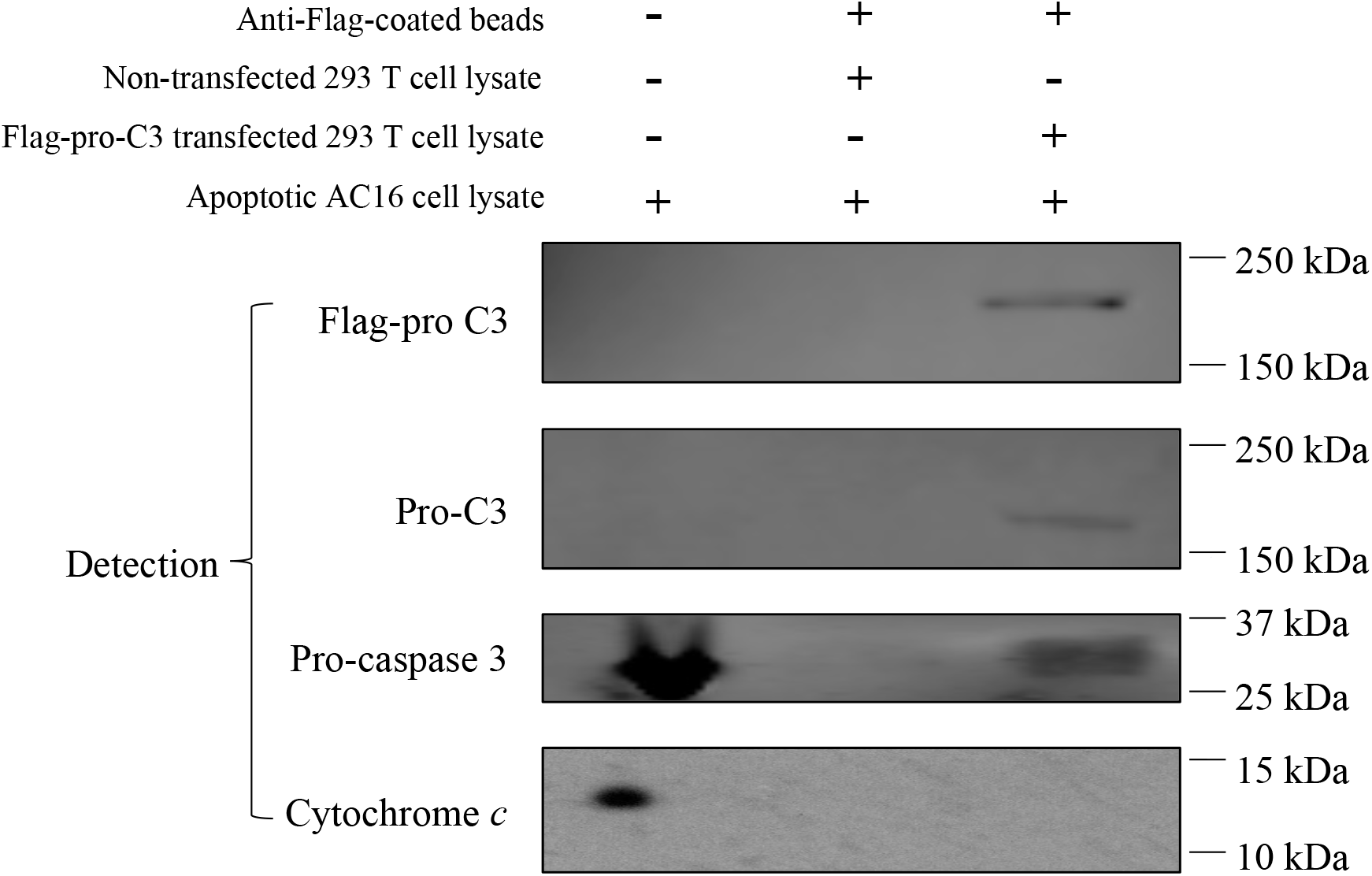
Pro-C3 binding with pro-caspase 3 in a cell-free system. Lane 1: AC16 cells were treated with 500mM H_2_O_2_ for 30 mins. Cell lysate was prepared as described in Methods. Lane 2: Anti-flag antibody coated beads were incubated with cell lysate of non-transfected 293 T cells (without Flag-pro-C3), then incubated with AC16 cell lysate. The pull-down materials were used for analyses. Lane 3: Anti-Flag antibody coated beads were incubated with cell lysate of transfected 293 T cells (contains Flag-pro-C3), then incubated with cell lysate of AC16 cells. The pull-down materials were then used for analyses.

## 4. Discussion

Our results showed that although human cardiomyocytes (AC16 cells) did not express C3, incubation of exogenous C3 reduced apoptosis in a cell culture system of cardiomyocytes (Fig.1) as well as in a cell-free apoptosis system (Fig. 2). Furthermore, pro-C3 was found to bind with an apoptotic factor, pro-caspase 3, in a cell-free system (Fig. 3). Thus, we presented firsthand evidence that exogenous C3 may interact with factor(s) in the intrinsic apoptotic pathway to inhibit apoptosis in cardiomyocytes.

Regarding how the exogenous C3 can interact with apoptotic factor(s) in cytosol, our pilot studies suggested that although AC16 cells did not produce matured C3, they were readily uptake the exogenous native C3 from extracellular milieu (Zhou, unpublished results). C3 uptake has been reported in human immune cells (Elvington et al., 2017), airway epithelial cells (Kulkarni et al., 2019), and retinal epithelial cells (Kaur et al., 2018). In particular, C3 uptake protects human airway epithelial cells from H_2_O_2_ induced cell death (Kulkarni et al., 2019).

Therefore, the anti-apoptotic function of C3 could be carried out by the exogenous C3 after its uptake by AC16 cells.

We recently found that Cyt *c* is present in the C3-binding complex in a murine myocardial I/R model which apoptosis is regulated (Fang et al., 2023). However, our current results suggest human C3 does not directly interact with Cyt *c* but rather downstream factor(s) in the intrinsic apoptosis pathway (Fig. 2). One explanation is that Cyt *c* may be present in the apoptotic complex in human cardiomyocyte to initiate the intrinsic apoptosis pathway, but the continuation of apoptosis process is blocked when C3 binds to a downstream factor in the pathway. An alternative explanation is that species differences may be manifested in the interactions between C3 and apoptotic factor(s) in mouse versus human.

We did attempt to use pull-down assay to identify the potential apoptotic factor(s) that C3 directly interact with. Our results showed that pro-C3 can bind with pro-caspase 3 (Fig. 3). While this finding is novel and interesting, it also opens some new scientific queries. For instance, pro-C3 is known as a single chain protein typically inside the cell before it being enzymatically processed into 2 chains linked by disulfate bonds. Traditionally pro-C3 is thought not functionally active, but our finding of pro-C3 binding with pro-caspase 3 suggests that it may function inside the cell by regulating apoptosis. Nevertheless, the purified C3 we used in anti-apoptotic experiments (Figs. 1, 2, and 3) were from human plasma and thus in the double-chain form. It remains to be determined if the double-chain form of C3 binds to specific factor(s) in the intrinsic apoptosis pathway similar to that of the single-chain form. Future research on this line may provide more insights.

Accumulative evidence indicates that the role of C3 in regulating apoptosis appears to be pertinent to specific cell types and pathological conditions. For instance, C3 exhibited pro-apoptotic effects in murine retina during photo-oxidative damage (Jiao et al., 2020) and in primate kidney during hemorrhagic shock (Halbgebauer et al., 2020). On the other hand, C3 was reported to also have anti-apoptotic function. For example, C3 can protect pancreatic β-cells from cytokine-induced apoptosis (Dos Santos et al., 2017) and possibly interacts with the autophagy-associated protein ATG16L1 (King et al., 2019a; King et al., 2019b). In addition, C3 seems to protect airway epithelial cells in a cigarette smoking model (Pei et al., 2022), and airway epithelial cells readily take up C3 from exogenous sources to mitigate apoptosis (Kulkarni et al., 2019). C3 was also reported to inhibit apoptosis in bullous-like skin inflammation (Zheng et al., 2020). Taking together with our current findings, C3 may act as a double-edged sword in regulating cell apoptosis depending on the pathologic conditions.

## Acknowledgements

The authors thank Drs. James Cottrell and David Wlody for their continued support. We also thank Drs. John P. Atkinson, Hrishikesh Kulkarni and Michelle Elvington (all at Washington University) for their scientific input and support in our pilot studies.

