## Supplemental Figures for "Complement C3 reduces apoptosis in human cardiomyocytes"

### Slide 1
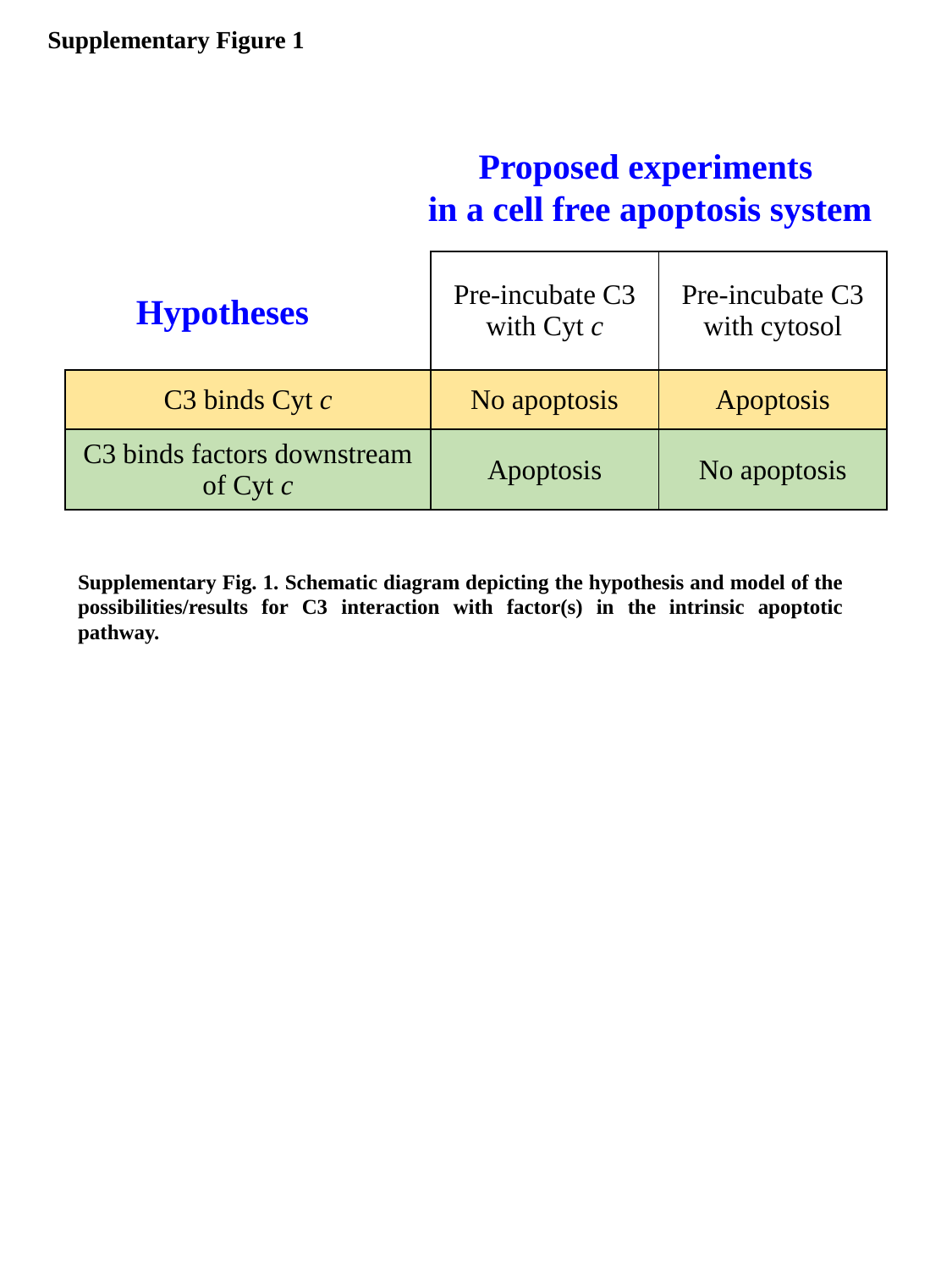

Supplementary Figure 1
Proposed experiments
in a cell free apoptosis system
| | Pre-incubate C3 with Cyt c | Pre-incubate C3 with cytosol |
| --- | --- | --- |
| C3 binds Cyt c | No apoptosis | Apoptosis |
| C3 binds factors downstream of Cyt c | Apoptosis | No apoptosis |
Hypotheses
Supplementary Fig. 1. Schematic diagram depicting the hypothesis and model of the possibilities/results for C3 interaction with factor(s) in the intrinsic apoptotic pathway.

### Slide 2
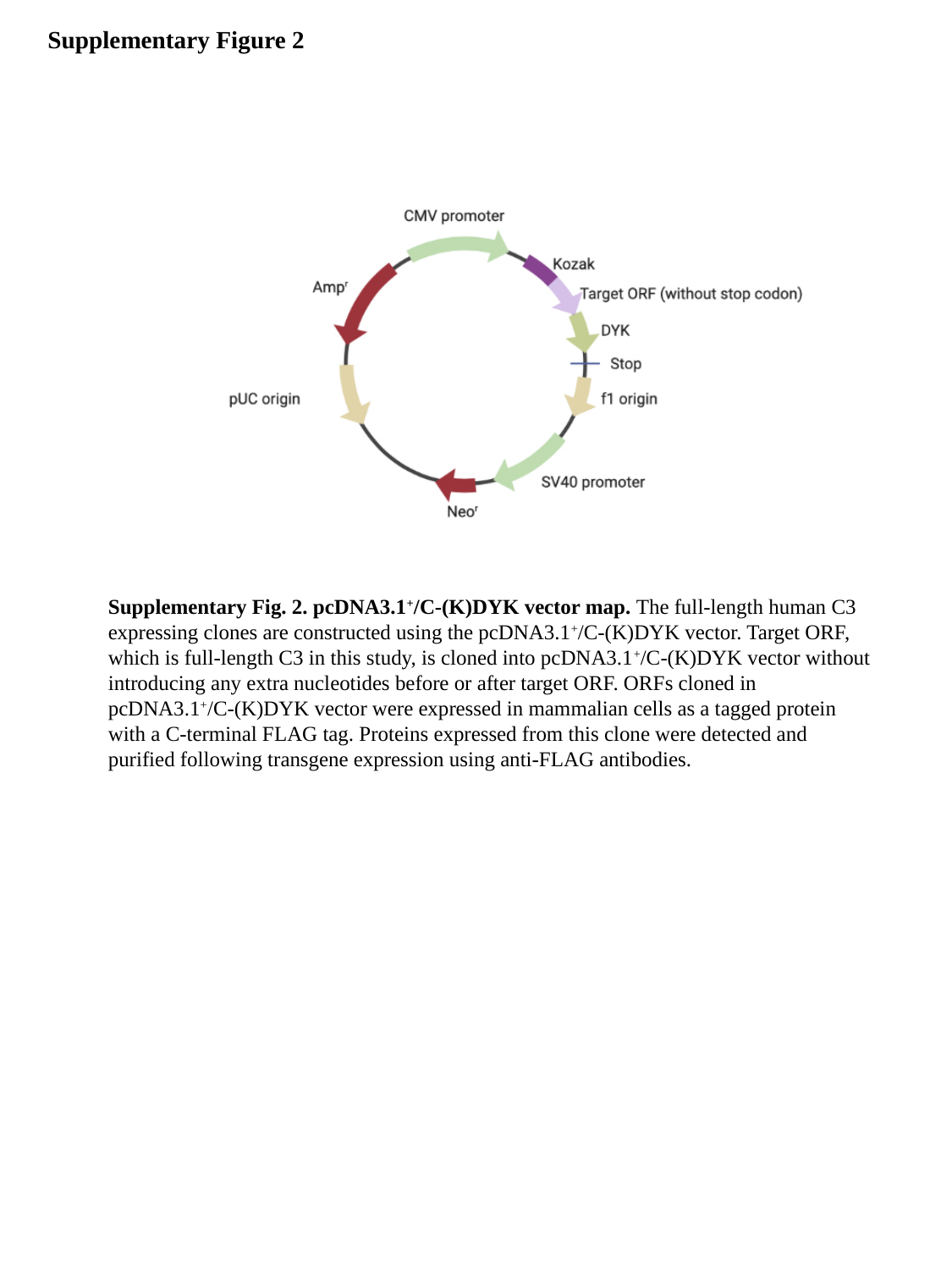

Supplementary Figure 2
Supplementary Fig. 2. pcDNA3.1^+^/C-(K)DYK vector map. The full-length human C3 expressing clones are constructed using the pcDNA3.1^+^/C-(K)DYK vector. Target ORF, which is full-length C3 in this study, is cloned into pcDNA3.1^+^/C-(K)DYK vector without introducing any extra nucleotides before or after target ORF. ORFs cloned in pcDNA3.1^+^/C-(K)DYK vector were expressed in mammalian cells as a tagged protein with a C-terminal FLAG tag. Proteins expressed from this clone were detected and purified following transgene expression using anti-FLAG antibodies.
